# Impaired response to mismatch novelty in the Li^2+^-pilocarpine rat model of TLE: correlation with hippocampal monoaminergic inputs

**DOI:** 10.1101/2024.02.07.579299

**Authors:** C Nascimento, V Guerreiro-Pinto, S Pawlak, A Caulino-Rocha, L Amat-Garcia, D Cunha-Reis

## Abstract

Novelty detection, crucial to episodic memory formation, is impaired in epileptic patients with mesial temporal lobe resection. Mismatch novelty detection, that activates the hippocampal CA1 area in humans and is vital for memory reformulation and reconsolidation, is also impaired in patients with hippocampal lesions. In this work we investigated the response to mismatch novelty, as occurs with the new location of known objects in a familiar environment, in the Li^2+^-pilocarpine rat model of TLE and its correlation with hippocampal monoaminergic markers. Animals showing spontaneous recurrent seizures (SRSs) for at least 4 weeks at the time of behavioural testing showed impaired spatial learning in the radial arm maze, as described. Concurrently, SRSs rats displayed impaired exploratory responses to mismatch novelty, yet novel object recognition was not significantly affected in SRSs rats. While the levels of serotonin and dopamine transporters were mildly decreased in hippocampal membranes from SRS rats, the levels on the norepinephrine transporter, tyrosine hydroxylase and dopamine-β-hydroxylase were enhanced, hinting for an augmentation, rather than an impairment in noradrenergic function in SRS animals. Altogether, this reveals that mismatch novelty detection is particularly affected by hippocampal damage associated to the Li^2+^-pilocarpine model of epilepsy 4-8 weeks after the onset of SRSs and suggests that deficits in mismatch novelty detection may substantially contribute to cognitive impairment in MTLE. As such, behavioural tasks based on these aspects of mismatch novelty may prove useful in the development of cognitive therapy strategies aiming to rescue cognitive deficits observed in epilepsy.

## Introduction

Epilepsy, a complex disease characterized by the development of recurrent, unprovoked seizures, affects over 65 million people worldwide and is one of the most common, chronic, serious neurological diseases and a *major* burden for public health systems [1–3]. Epilepsy is frequently allied with neurological comorbidities, such as cognitive deficits, depression, anxiety and psychiatric disturbances [1,3,4]. Mesial temporal lobe epilepsy one of the most prevalent forms of epilepsy, is characterized by mesial temporal lobe symptoms affecting the hippocampus, parahippocampal gyrus and amygdala, and is frequently (in 56% of cases) linked to hippocampal sclerosis (MTLE-HS) [5]. Impaired cognitive functioning including language, executive function and declarative memory deficits [6] are a major hallmark of MTLE-HS. A few studies have also reported that MTLE patients present attention deficits and arousal abnormalities, that can greatly affect novelty processing not only by the dorsal attention network but also through the mesial temporal lobe, affecting particularly hippocampal-dependent tasks [7–10]. Since only about 11-26% of patients with MTLE-HS achieve complete seizure control under pharmacological treatment with existing multiple antiseizure drugs (ASDs), finding new therapeutic strategies to prevent progressive cognitive decline in epilepsy patients is a main priority.

Novelty is an important stimulus in episodic memory formation, different aspects of novelty distinctively impact hippocampal dependent learning and synaptic plasticity. In fact, hippocampal long-term potentiation (LTP) or long-term depression (LTD) of hippocampal synaptic transmission, encode different aspects of novelty acquisition [11], LTD being facilitated during the location of new objects or known objects in new locations, a behavioural mismatch novelty paradigm, and LTP being favoured during exploration of a new environment, key to the formation of spatial maps and memory consolidation [11–14]. Memories and associated synaptic plasticity processes, are also shaped by previous learning experiences (either recent or remote) through metaplasticity [15]. Novelty is an important trigger for these metaplastic changes. Furthermore, while spatial novelty mainly enhances retrieval of a previously acquired memory [16], mismatch novelty paradigms were shown to enhance inhibitory avoidance learning, by influencing hippocampal LTD [13].

Interestingly, mismatch novelty detection, particularly important in memory reformulation and reconsolidation, has also been associated with the activation of the hippocampal CA1 area in human studies, and is impaired in patients with hippocampal lesions [17,18]. Furthermore, studies from our group showed that repeated exposure to mismatch novelty increases both LTP and LTD in the hippocampus [19], suggesting that behavioural tasks involving mismatch novelty features may reveal useful in the development of cognitive therapy strategies aiming to reduce the LTP/LTD imbalance as found in aging or diseases like epilepsy or Dowńs syndrome.

Several neurotransmitter systems have been implicated in the hippocampal processing and regulation of novelty stimuli. Among these, monoaminergic projections from the *ventral tegmental area* (VTA), *locus coeruleus* and *median raphe* to the hippocampus have been implicated in regulating physiological arousal, attention, and motivation and are thought to play an essential role in the efficiency of cognitive function [20–23] while playing a crucial role in recognition memory and novelty signalling [23]. Altered monoaminergic neurotransmission not only constitutes a risk factor for the development of epilepsy [24] but is either linked to the degeneration or upregulation of ascending projections to the hippocampus and cortex, altered neurotransmitter levels or altered monoamine receptor levels and function has been reported in the hippocampus of MTLE patients [25]. Similar observations occurred in experimental models of epilepsy, including loss of limbic-projecting serotonergic neurons from the *median raphe nucleus* [26], and deterioration of dopaminergic projections from the VTA to the *nucleus accumbens* [27] coupled to decreased vesicular monoamine transporter 2 in the temporal cortex and hippocampus [28].

In this work we investigated the exploratory response to mismatch novelty in the Li^2+^-pilocarpine rat model of TLE and characterized hippocampal monoaminergic and synaptic markers while probing also other hippocampal dependent learning and memory tasks. Exploration of a known environment containing familiar objects presented in a new location was impaired in rats showing spontaneous recurrent seizures (SRSs) for at least 4 weeks, suggesting that deficits in mismatch novelty detection indeed contribute to cognitive impairment in MTLE. This was correlated with alterations in the hippocampal monoaminergic system that may contribute to the attention deficit-like profile previously observed in the Li^2+^-pilocarpine model of epilepsy.

## Materials and Methods

### Animals and induction of SRSs

Animals were housed in the local Animal House of the Institute of Physiology, Faculty of Medicine, University of Lisbon until use, and were maintained under a 12:12-h light/dark cycle at a temperature of 22°C, with food and water ad libitum. All procedures were in accordance with the standards established in the Guide for Care and Use of Laboratory Animals, the Portuguese and European law on animal welfare and were approved by the Ethical Committee of the Faculty of Medicine, University of Lisbon.

Adult (12-week-old, 335-375g, n=32) male Wistar rats were handled twice a day for three days and status epilepticus (***SE***) was induced (n=24) by intraperitoneal pilocarpine administration (10mg/Kg) essentially as described [29]. Animals were pre-treated with LiCl (300 mg/kg, i.p., Sigma) and twenty-four hours later, they received methyl-scopolamine (1 mg/kg, i.p., Sigma) to block the peripheral cholinergic effects of pilocarpine. ***SE*** was induced 15 min later by administering pilocarpine (10mg/Kg) and, if necessary, additional lower doses (5 mg/Kg) of pilocarpine were administered every 20 min until either ***SE*** is observed or a maximum of four doses was attained. Behavioural epileptiform seizures were monitored for 30min, scored according to the scale of Racine modified by Lüttjohann *et al*. (2009) [30] and terminated by diazepam (i.p., 10 mg/Kg) delivery. Xilazine (i.m., 10 mg/Kg) was administered immediately after seizure onset to prevent muscle exhaustion caused by convulsions. Seizure recurrence within the next 24-48h was controlled, when required, with additional diazepam (i.p., 5-10 mg/Kg). This greatly increased animal survival. ***Sham*** animals (n=24) were subjected to the same procedure except for the administration of pilocarpine that was replaced by an equivalent volume of saline (NaCl, 0.9%). Occurrence of spontaneous recurrent seizures (***SRSs***) was detected by 24h video monitoring within 4-10 weeks of pilocarpine administration and the frequency and severity of behavioural seizures were scored as described [30]. Behaviour was evaluated 4-12 weeks following the detection of the first unprovoked seizure (6-10 months of age). Animals presenting SRSs (n=24) and respective Sham controls (n=24) were subjected to a global evaluation of their motor capacity and anxiety levels using the elevated plus maze (***EPM***) test and the open-field (***OF***) test. Learning and memory impairment was evaluated in a subset of animals using the 8-arm radial arm maze (***RAM***, n=8 Sham/SRSs) test for spatial memory or the novel object recognition (***NOR***, n=8 Sham/SRSs) test for non-spatial memory. Evaluation of the mismatch novelty (***MN***, n=16 Sham/SRSs) response using the holeboard with objects was performed as described with minor modifications [11] starting 24h after the ***OF***. All behavioural testing/training sessions were performed between 9:00 a.m. and 17:00 p.m. in a sound attenuated room. Each trial was video recorded and analysed using the video-tracking software ANY-maze (Stoelting, Europe).

### Evaluation of anxiety and locomotion using the EPM and OF

Individual levels of anxiety and general locomotor behaviour, was evaluated using the OF test [31] and the EPM test as adapted by Schneider [32] essentially as previously described [19].

The Elevated plus-maze (EPM) consisted of two open arms (50cmx10cm) and two enclosed arms (50cmx10cmx40cm), extending from a central platform (10cmx10cm) and raised 50cm above floor level. During the EPM test, the animal was placed on the central platform, facing the open arm, and allowed to explore the maze for 5 minutes. The maze was cleaned with a 70% ethanol solution between each animal trial. Animal activity was either monitored manually by the experimenter or video recorded and analysed using the video-tracking software Anymaze. The parameters scored were the number of entries in open/closed arms, the time spent in open/closed arms, the time on the central platform, the distance travelled by the animal in the entire maze [32], and the number of rearings (considered when the rat was on its hind legs, touching or not with his front paws on the wall).

The Open Field (OF) test consisted of the exploration of a large square chamber (66cmx66cm wide, 60cm-high walls), for 5 minutes. The apparatus was divided into three virtual zones (a central square 20×20cm, an intermediate zone and a peripheral zone 15cm-wide adjacent to the walls) for analysis of the rat behaviour. Animals were removed from the home cage and placed directly in the centre of the apparatus. The movement of the animal in the arena during the test session was recorded and the performance of the animals was evaluated by the escape latency (s), total distance travelled, the number of rearings, the number of entries and the time spent in each virtual zone [31].

### Evaluation of cognitive performance in the RAM

The cognitive performance of Sham versus SRSs rats was evaluated using the radial arm maze (RAM) test. The RAM was first used to evaluate the ability of rats to memorize the location of baited arms upon one-week repeated exposure to the baited RAM. The RAM consisted in an octagonal centre platform 27cm in diameter connected to eight equally spaced arms, each measuring 50cmx10cm and 20cm-higth, with a cylindrical food cup (3.5cm ø, 0.5cm depth) at the end of each arm. The maze was elevated 50cm from the floor and was surrounded by several extra-maze cues. The parameters evaluated in the RAM were arm entries, counted if all four paws were placed on that arm (He, Yamada, Nakajima, Kamei, & Nabeshima, 2002); arm latencies (maximum time to find the three available food rewards) and the number of rearings. Arm entries were scored as errors of working memory (re-entries into baited arms) and errors of reference memory (entries into non-baited arms).

Before each trial animals were food deprived (maintained at 85–90% of free feeding body weight) (Crusio & Schwegler, 2005). Rats were placed in the centre of the apparatus At the beginning of the test, surrounded by an opaque cylinder, that was kept for 5 seconds before the animal was allowed to perform arms choices. Spatial cues were placed in the room walls to allow animals to locate the three food rewards, always placed in the same set of three arms. To prevent animals from using within-maze cues, the maze was rotated 45° at the end of each day (between subsequent trials), so that intra-maze and extra-maze cues were dissociated. The test duration was 10 min on the first trial (maze recognition) and 5 min on the remaining trials (2 trials per day). Cognitive performance was evaluated by day by averaging performance in the two trials.

### Novel Object Recognition (NOR)

The NOR test was performed in a square arena essentially as described [33] and was composed of three sessions: habituation, training, and test sessions [23]. The habituation session consisted of one session of free exploration in the arena (5 min). In the training session, two objects (similar in size and shape but different in texture, colour, and patterning) were added to the arena (Fig. 2), and this location was kept for the persisting object in the subsequent test session. Animals were placed in the middle of the arena facing away from the objects and allowed to freely move for 5 min. Test for novel object recognition was performed 24h latter by replacing one of the previously experienced objects (familiar objects F1 and F2) by a novel object (N). Exploration was scored when the animal touched an object with its forepaws or snout, bit, licked, or sniffed the object from no more than 1.5 cm. Animal movements and the time spent exploring each object were analysed. Animals spending less than 20s exploring the objects were excluded from the study. Exploration of novel objects was scored using: 1) the object preference index - ratio between the time spent exploring one object over the total time spent exploring both objects [t_F1_ or t_F2_/(t_F1_ + t_F2_)]; 2) the object recognition index - ratio between the time spent exploring the novel object and the total time spent exploring both objects [t_N_/(t_N_ + t_F_)], an index of memory retention; and 3) the object discrimination index - ratio between the difference between the time spent exploring the novel and the familiar object, and the total time spent exploring both objects [(t_N_ – t_F_)/(t_N_ + t_F_)], that allowed visualization of data with no memory retention scored as zero.

### Mismatch novelty test

The mismatch novelty (MN) test consisted in the exploration of the novel location of known objects in a familiar environment essentially as described (Supp. Fig. 1) [13]. This consisted of a holeboard composed by an arena (66×66cm, 60cm-high walls) and containing one hole at each corner (four holes, 5.5cm diameter, 4.5cm deep). Animal habituation and testing was performed in 5-min sessions. One day prior to object exposure, all animals were exposed to the empty holeboard to get accustomed to the environment. Objects were introduced on the next day (1^st^ exposure) in three of the four holes for all animals. On the second day animals were either exposed to the same spatial distribution of objects (re-exposure) or to a new spatial configuration of the objects (novel configuration). Each trial was video monitored, recorded, and later analysed using automated video-tracking software (Anymaze software, Stoelting, Europe). The parameters scored were the travelled distance, the number of entries, and the time spent in each virtual zone of the apparatus (central, intermediate, and peripheral zones; essentially as defined for the OF). Exploration of objects and general exploratory activity were evaluated by the number of nose pokes and the number of rearings, respectively. Increased nose pokes vs rearings were taken as a positive response to novelty.

### Western blot analysis of monoaminergic markers and synaptic proteins

For western blot studies total hippocampal membranes were isolated essentially as previously described [34]. Briefly, the hippocampi of Sham and SRSs rats were dissected and collected in sucrose solution (320mM Sucrose, 1mg/ml BSA, 10mM HEPES e 1mM EDTA, pH 7,4) containing protease (complete, mini, EDTA-free Protease Inhibitor Cocktail, Sigma) and phosphatase (1 mM PMSF, 2 mM Na3VO4, and 10 mM NaF) inhibitors, homogenized with a Potter-Elvejham apparatus and centrifuged at 1500g for 10 min. The supernatant was collected and further centrifuged at 14000g for 12 min. The pellet was washed twice with modified aCSF (20mM HEPES, 1mM MgCl2, 1.2mM NaH2PO4, 2.7mM NaCl; 3mM KCl, 1.2mM CaCl2, 10mM glucose, pH 7.4) also containing protease and phosphatase inhibitors and resuspended in modified aCSF to a concentration of 1mg/ml protein. Aliquots of this suspension of hippocampal membranes were snap-frozen in liquid nitrogen and stored at -80°C until use.

For western blot, samples incubated at 95°C for 5 min with Laemmli buffer (125mM Tris-BASE, 4% SDS, 50% glycerol, 0,02% Bromophenol Blue, 10% β-mercaptoethanol), were run on standard 10% sodium dodecyl sulphate polyacrylamide gel electrophoresis (SDS-PAGE) and transferred to PVDF membranes (Immobilon-P transfer membrane PVDF, pore size 0.45 μm, Immobilon). These were then blocked for 1 h with either 3% BSA or 5% milk solution in Tris-buffered saline containing 1% Tween (TBST) and incubated overnight at 4°C with rabbit anti dopamine transporter (DAT, 1:2000, Proteintech Cat# 22524-1-AP, RRID:AB_2879116), mouse monoclonal anti norepinephrine transporter (NET, 1:500, Atlas Antibodies Cat# AMAb91116, RRID:AB_2665806), rabbit anti serotonin transporter (SERT, 1:1000, Proteintech Cat# 19559-1-AP; AB_2878590), rabbit anti-tyrosine hydroxylase (TH, 1:1000, Abcam Cat# Ab112; RRID:AB_297840), mouse monoclonal anti-gephyrin (1:3000, Synaptic Systems #147011, RRID:AB_2810215), rabbit anti-PSD-95 (1:750, Cell Signalling Technology #2507, RRID:AB_561221), rabbit anti-GluA1 (1:4000, Millipore Cat# AB1504; RRID:AB_2113602), rabbit anti-GluA2 (1:1000, Proteintech Cat# 11994-1-AP; RRID: AB_2113725), mouse monoclonal anti-GluN1 (1:1000, Proteintech Cat# 67717-1-Ig; RRID: AB_2882906), rabbit anti GluN2B (1:1000, Cell Signalling Technology Cat#4207; RRID: AB_1264223), rabbit anti-synaptophysin (1:7500, Synaptic Systems #101002, RRID:AB_887905), and either mouse monoclonal anti-β-actin (1:5000, Proteintech #60008-1-Ig, RRID: AB_2289225) or rabbit polyclonal anti-α-tubulin (1:4000, Proteintech #PT11224-1-AP, RRID: AB_ 2210206) primary antibodies. After washing 3x for 10 min with TBST, the membranes were incubated for 1h with anti-rabbit IgG or anti-mouse IgG secondary antibody both conjugated with horseradish peroxidase (HRP) (Proteintech) at room temperature. Excess bound secondary antibody was then removed by washing and HRP activity was visualized by enhanced chemiluminescence with Clarity ECL Western Blotting Detection System (Bio-Rad). Band intensity was evaluated with the Image J software using either β-actin or α-tubulin band density as loading control. Immunostaining of the different targets was normalized to loading control band density and differences in target protein expression in SRS animals were expressed as percentage change relative to Sham controls.

### Statistics

Values of behavioural assessment parameters are presented as the mean ± S.E.M of 6-20 animals. In western blot experiments each *n* represents one experiment in a single animal. Significance of the differences between the Sham and SRSs groups was calculated by Student’s t test with Welch correction for unequal variances. Differences were considered significant for P values of 0.05 or less. Statistical analysis was performed using GraphPad Prism 6.01 for Windows.

## Results

### Elevated Plus Maze (EPM) and Open-field (OF) tests

During rodent exploration of the EPM (Fig. 1.A-E) the time spent by Sham animals in the open arms (5.4±1.6%, n=24) was much lower than the time spent in the closed arms (78.8±3.2%, n=24). In the remaining time (15.8±3.9%, n=24) the animals were in the centre of apparatus (the crossing of closed and open arms). In contrast, SRSs animals spent more time in the open arms (30.5±5.8%, n=24) and less time in the closed arms open arms (55.2±6.0%, n=24). Accordingly, Sham animals entered the open arms less (1.2±0.3, n=24) than SRSs rats (3.5±0.7%, n=24), and the opposite was observed with the number of entries in the closed arms (4.9±0.5%, n=24 for Sham vs. 6.5±0.8, n=24 for SRSs animals) from this central position. The total distance travelled in the EPM during the 5-min trial was larger for SRSs (0.92±0.08m, n=24) than for Sham rats (0.62±0.5m, n=24). Rearings were almost entirely performed within the closed arms and were slightly higher for SRSs (10.0±0.9, n=24) when compared to Sham rats (8.8±0.7, n=24).

**Figure 1.**
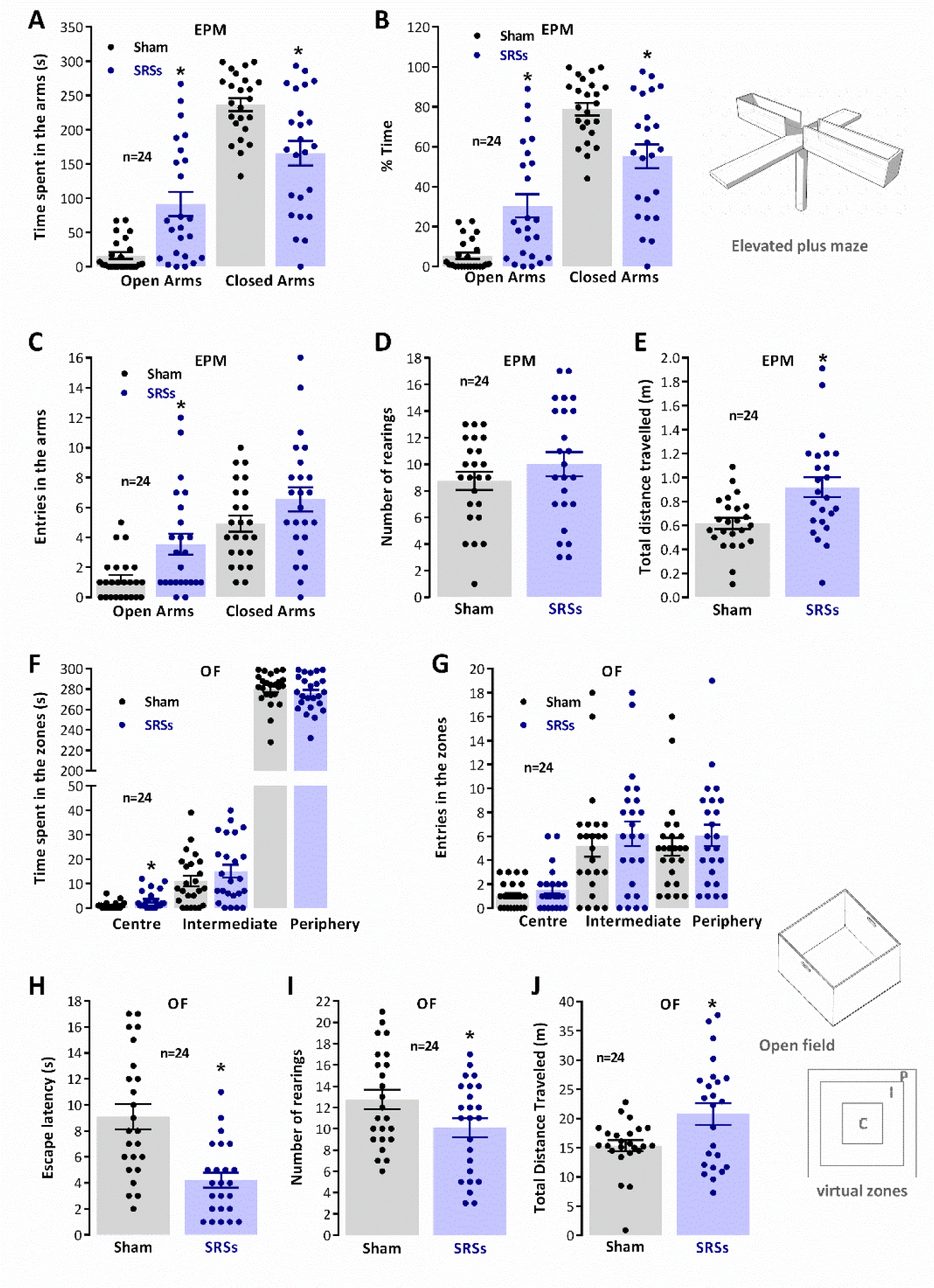
– Enhanced risk-taking behaviour and locomotion in the Li^2+^-pilocarpine rat model of epilepsy. Perception of risk by SRSs rats was evaluated in the EPM as given by the time spent in the open and closed arms (**A.-B.**) and number of entries in the arms (**C.**). A schematic representation of the apparatus is depicted (top, right). The number of rearings performed during the trial (**D.**) and the total distance travelled in the apparatus (**E.**) were used to evaluate total EPM exploratory behaviour. Thigmotaxis in the OF was evaluated by measuring the time spent in the different virtual zones of the OF (**F.**) and the number of interzone crossings (**G.**), as defined in the schematic representation of the OF (**G.**, left). Escape latency (**H.**), time for animals to begin locomotion and trial start, was taken as a measure of impulsive behaviour. Global exploratory activity in the OF was evaluated by the number of rearings (**I.**) and the total distance travelled (**J.**). Values are the mean ± S.E.M. Total trial duration was of 5 min for each EPM or OF session. *p < 0.05, (One-way ANOVA) vs Sham controls, followed by Sidak’s multiple comparison test for time spent and number of entries in the arms in RAM and time and number of entries in the zones of the OF. For the number of rearings, total distance travelled, and escape latencies the student’s t test was used instead.

Behaviour in the OF during the 5-min trial (Fig. 1.F-J) was analysed through the definition of three virtual zones (peripheral, intermediate and central zones). Sham animals showed a high thigmotaxis (Fig. 1.A) as evidenced by the minimum time spent in the centre (0.9±0.3s, n=24) and intermediate (11.0±2.1s, n=24) zones of the OF and preference for remaining in the periphery zone (286.4±3.4s, n=24), as previously described for naïve animals of similar age. Comparatively, the time spent by SRSs animals in the central (3.0±0.8s, n=24) zone was higher than for Sham animals while the time spent in the intermediate (15.1±2.7s, n=24) and periphery (280.9±3.6s, n=24) zones did not substantially different from Sham rats. The global locomotor activity, accessed by the total distance travelled in the open-field, was higher in SRSs animals (20.8±1.8m, n=24) as compared to the one covered by Sham rats (15.4±0.9m, n=24). Sham and SRSs animals did not present significant differences in the number of entries in any of the different zones. The total number of rearings was lower for SRSs animals (10.1±0.9, n=24) as compared to Sham animals (12.8±0.9, n=24), and reflects rearings performed predominantly in the periphery zone.

### Spatial learning in the RAM

During exploration of the RAM the latency to find the 1^st^ baited arm Supp. Fig. 2.A) was not significantly different between Sham and SRS animals during the whole testing period. However, the latency to find the 2^nd^ and 3^rd^ baited arms was higher in SRSs rats (Supp. Fig. 2.B and C). The latency to task completion was significantly higher from the third day of testing (270.1±19.9s, n=8 for SRS vs 181.0±16.4s, n=8 for Sham on day 3) and SRSs rats showed little improvement in their performance from the 1st to the 5th day of testing (Supp. Fig. 2.C), and three of the SRS animals were unable to complete the task on day 5, evidencing a markedly decreased capacity to memorize the location of the baited arms. The distance travelled in the RAM (Supp. Fig. 2.D) was higher on the first day for SRSs rats (30.9±2.8, n=8 for SRS vs 23.7±1.4, n=8 for Sham) but no significant differences between the two groups were encountered in the remaining days. The number of rearings performed during the RAM training and test sessions (Supp. Fig. 2.E) were higher for SRS rats on the first day (33.9±2.5, n=8 for SRS vs 27.3±2.4, n=8 for Sham) but decreased markedly throughout the sessions being much lower that the ones performed by Sham rats by the last day of the RAM test (11.5±1.3, n=8 for SRS vs 19.0±1.9, n=8 for Sham).

### Novel object recognition

The NOR task consisted of a training session when animals were exposed to previously unknown objects and a test session, delivered 24 h later, when one of these two now familiar object and one novel object were presented. During the training session, we observed that Sham animals explored both objects equally (Fig. 2.A, ***Obj. 1***: 51.2 ± 2.4%, ***Obj. 2***: 48.8 ± 2.4%; n = 8), as given by the object preference index (Fig. 2.G), whereas during the test session, the animals explored the novel object to a significantly larger extent than the familiar object (Fig. 2.D, ***F Obj.***: 61.5 ± 1.5%, ***N Obj.***: 38.5 ± 1.5%; n = 8). This is a clear behavioural indication that Sham rats successfully formed a memory of the objects lasting for 24h during the training phase. SRSs animals showed a similar behaviour during both the training session (Fig. 2.A, ***Obj. 1***: 48.4 ± 3.5%, ***Obj. 2***: 51.6 ± 3.5%; n = 8, Fig. 2.G) or when a novel object was introduced 24h later (Fig. 2.D, ***F Obj.***: 59.9 ± 0.8%, ***N Obj.***: 40.1 ± 0.8%; n = 8), suggesting that novel object recognition was not significantly affected in SRS rats. This was also evidenced by absence of differences in the object recognition and discrimination indexes (Fig. 2H, 3I), despite the marked differences observed in general exploration between Sham and SRSs animals as given by the number of rearings during the training (7.0±1.0, n=8 for SRS vs 15.8±1.5, n=8 for Sham, Fig. 2B) and the test sessions (5.9±1.0, n=8 for SRS vs 14.5±1.7, n=8 for Sham, Fig. 2E) and by the total distance travelled in the training (3.9±0.4m, n=8 for SRS vs 1.8±0.3m, n=8 for Sham, Fig. 2C) and test session (2.9±0.3m, n=8 for SRS vs 0.9±0.2m, n=8 for Sham, Fig. 2F).

### Mismatch novelty detection

The MN test consisted of the exploration of the novel location of known objects in a holeboard in a familiar environment essentially as described (cite). In the first session animals were exposed to the empty holeboard and in the following session objects were introduced. In the third session Sham and SRS animals were divided into two groups one being exposed to the same object configuration and other exposed to the novel object configuration (Fig. 3.A). The behavioural responses of Sham and SRSs animals were monitored for 5 min.

**Figure 3.**
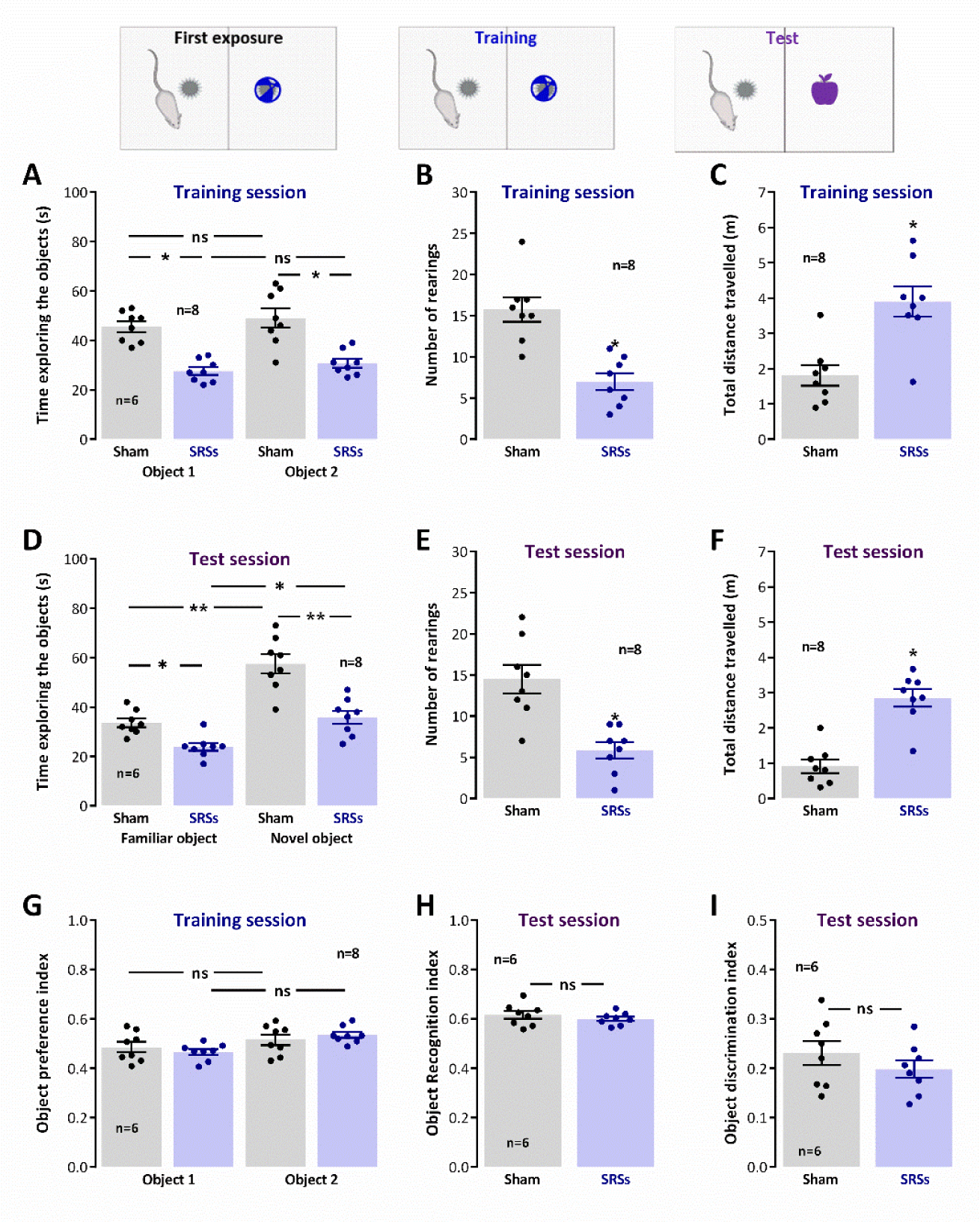
– Capacity for novel object recognition is not significantly affected in the Li^2+^-pilocarpine rat model of epilepsy. Exploration of the objects during the training and test sessions was evaluated by accessing the total time exploring the objects (**A., D.**) assessed by the time spent in close vicinity (sniffing/facing the object from <1.5 cm) or contacting the object with the forepaws or snout and biting or licking the object. Global exploratory activity in the arena was evaluated by the number of rearings (**B., E.**) and total distance travelled (**C., F.**) during each trial. Novel object exploration was evaluated by the object preference index (**G.**) in training sessions and by the object recognition (**H.**) and object discrimination (**I.**) indexes in the test sessions. Total trial duration was of 5 min for all sessions. Values are the mean ± S.E.M. *p < 0.05 (Student’s t-test) vs Sham animals.

Upon the first exposure to the holeboard, the general exploratory activity of Sham animals was lower than the one observed in the OF maze, as evidenced by the decrease in the number of rearings that was of 6.0±0.8 (n=16) in holeboard exploration and 12.8±0.9 (n=24) in the OF test, and by the decrease in the total distance travelled on the holeboard 13.0±1.4m (n=16) compared to the distance travelled in the open-field 15.4±0.9m (n=24). The animals spent slightly more time in the periphery zone of the holeboard (291.6±2.5s, n=16), where the holes are located, than in the periphery of the open-field (286.4±3.4s, n=24). Thus, animals although likely now less fearful, spent more time in the periphery exploring the holes, as evidenced by the number of nose-pokes (9.0±0.7, n=16). SRSs animals behaved similarly when exposed to the holeboard, since the total number of rearings (8.6±1.0, n=16) was only a bit higher for SRSs animals than the one performed by Sham animals, while the number of nose-pokes was slightly lower (7.8±1.2, n=16), yet the total distance travelled by SRSs animals (26.1±2.7m, n=16) in the holeboard was double the one travelled by Sham animals (13.0±1.4m, n=16), as occurring in virtually all other tests, a reflection of their anxious and attention deficit -like behaviour, as previously described [35,36].

Introduction of objects on the second day increased the number of nose-pokes for both Sham (15.3±1.0, n=16) and SRSs (10.6±1.0, n=16) animals (Fig. 3.C) but this effect was more patent for Sham animals since now there was a significant difference between Sham and SRSs behaviour. No noticeable differences were observed in the behaviour of Sham and SRSs animals regarding the number of rearings (Fig. 3.B), time spent (Fig. 3.E) or number of entries (Fig. 3.I) in the different zones of the apparatus in the session when the objects were first presented. Total distance travelled was nevertheless higher (P<0.05, t-test) for SRSs (20.4±2.1m, n=16) than for Sham animals (10.7±1.5m, n=16).

For the subgroup of animals subjected to re-exposure to the same configuration of objects, no significant differences (P>0.05) between Sham (n=8) and SRSs (n=8) animals in any of the behavioural parameters monitored (Fig. 3.B, C, F and J), including the total distance travelled (15.4±1.4m, n=8 for Sham vs. 16.5±3.3m, n=8 for SRSs). Conversely, for the subgroup exposed to a novel configuration of the objects, marked differences were observed between Sham and SRSs animals in global exploratory activity as given by the number of rearings (10.3±1.3m, n=8 for Sham vs. 6.8±1.5m, n=8 for SRSs) and total distance travelled (13.7±2.2m, n=8 for Sham vs. 28.6±3.3m, n=8 for SRSs). In addition, the number of nose pokes, a major indicator of a response to novelty, was also significantly higher for Sham (13.2±1.4, n=8, P<0.05) than for SRS animals (8.4±1.4, n=8), and significantly higher (P<0.05) than the one performed by Sham animals undergoing re-exposure to the same object arrangement (8.0±1.2, n=8). Altogether, this suggests an impairment in the response to mismatch novelty in SRSs rats.

### Evaluation of hippocampal levels of synaptic and monoaminergic markers

Numerous studies have identified marked differences in synaptic structure and molecular composition in epileptic rodent models. However, findings are often contradictory between different rodent models or even distinct rat strains. As such we briefly characterized the levels of a general synaptic marker (synaptophysin) and markers of both glutamatergic (PSD-95) and GABAergic synapses (gephyrin). The global levels of synaptophysin were not changed in hippocampal membranes of SRS rats as compared to Sham controls (n=6, P<0.05, Fig 4.C). Yet, both PSD-95 and gephyrin were significantly reduced to 82.7±5.0% (n=6, P<0.05, Fig 4.A) and 63.0±6.0% (n=6, P<0.05, Fig 4.B) of the levels observed in Sham animals.

**Figure 4.**
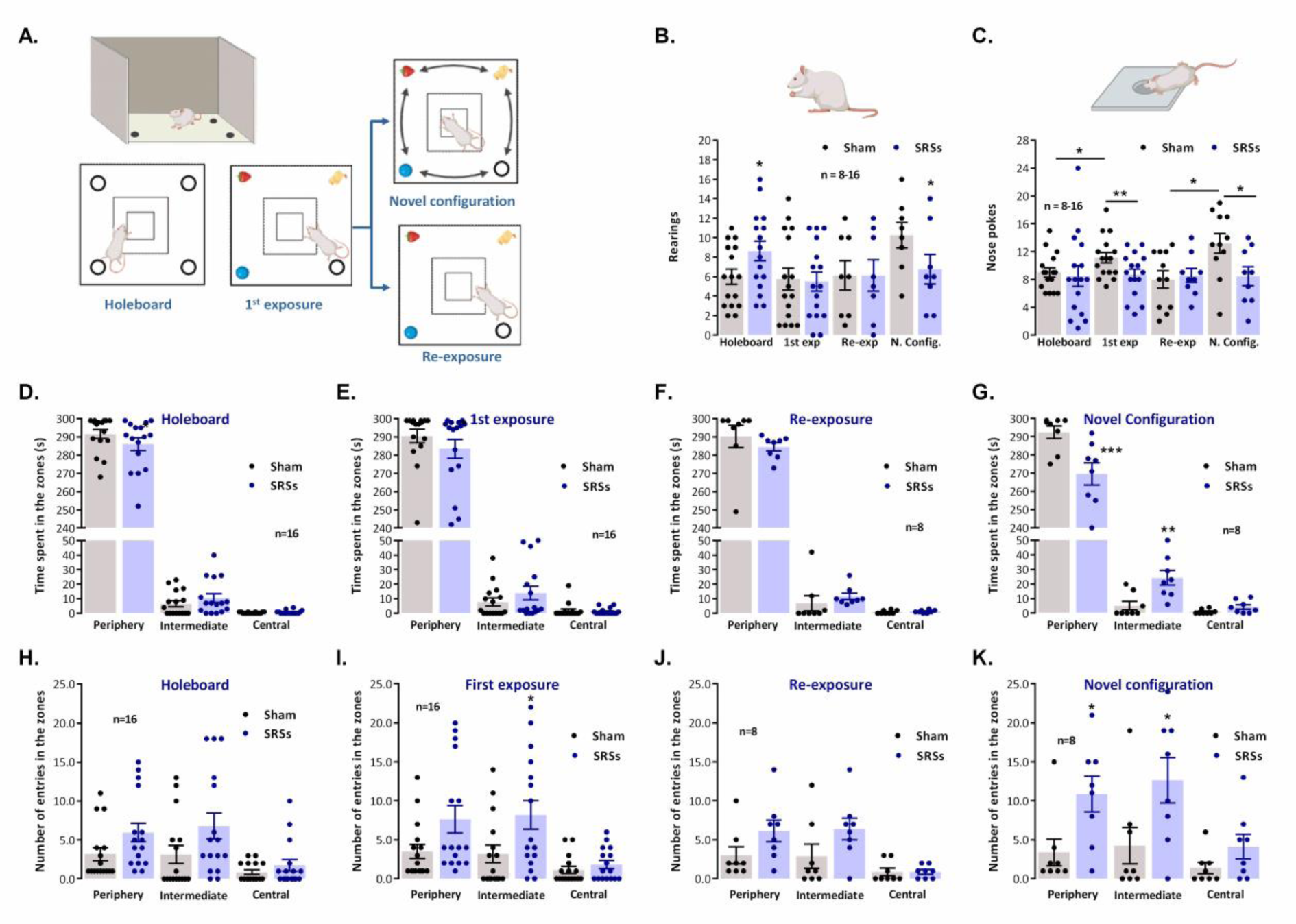
– Hippocampal synaptic composition is altered in the Li^2+^-pilocarpine rat model of epilepsy. Each panel shows at the bottom the western-blot immunodetection of PSD-95 (**A.**), gephyrin (**B.**) and synaptophysin-1 (**C.**) obtained in one individual experiment. Western blot experiments were performed using total hippocampal membranes isolated from five individual animals for both Sham and SRSs. Respective average change in PSD-95 (**A.**), gephyrin (**B.**), and synaptophysin-1 (**C.**) immunoreactivities are also plotted at the top in each panel. Individual values and the mean ± S.E.M of 5 independent experiments are depicted. 100% - averaged target protein immunoreactivity in Sham controls. **\*** P < 0.05 (Student’s t-test) as compared to Sham controls.

**Figure 4.**
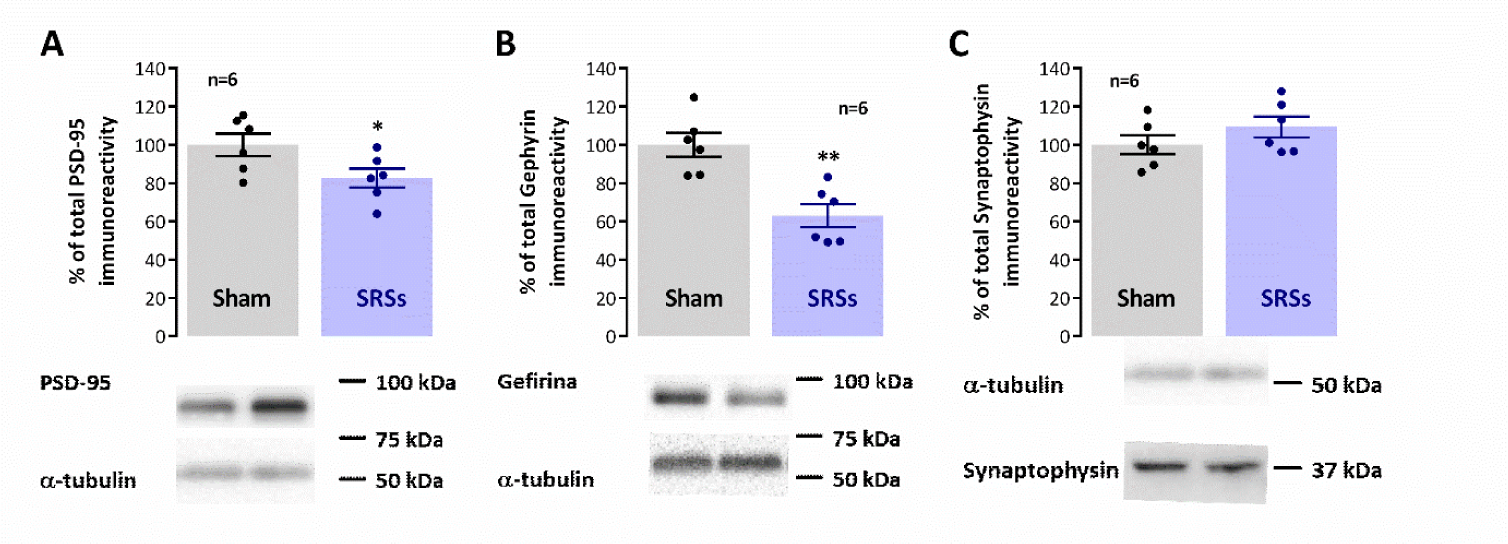
– Mismatch novelty response is impaired in the Li^2+^-pilocarpine rat model of epilepsy. A. Schematic representation of the holeboard apparatus and of the sequence of novelty test sessions. Rearings (**B.**) and nose-pokes (**C.**, head dips) in each of the four holes of the holeboard apparatus for the duration of the testing procedure for (from left to right) holeboard exposure (n=16), first exposure to the objects (n=16), re-exposure to the same configuration of objects (n=8) and exposure to a novel spatial configuration of the objects (n=8). Total distance travelled in the different virtual zones of the holeboard apparatus is shown during holeboard exposure (**D.**), first exposure to the objects (**E.**), re-exposure to the same configuration of objects (**F.**) and exposure to a novel spatial configuration of the objects (**G.**). Number of entries in the different virtual zones of the holeboard apparatus is shown during holeboard exposure (**H.**), first exposure to the objects (**I.**), re-exposure to the same configuration of objects (**J.**) and exposure to a novel spatial configuration of the objects (**K.**).Total duration of each daily training session was 5 min. Values are the mean ± S.E.M. *p < 0.05, **p < 0.01 (students t-test) as compared with sham animals for the same testing procedure.

Impaired synaptic plasticity, as observed in animal models of epilepsy is believed to be associated with cognitive deficits in epileptic patients. Changes in both NMDA and AMPA receptor subunit composition have also been demonstrated in animal models of epilepsy and are believed to contribute to altered cognitive ability human epilepsy. As such we also verified in our model how AMPA GluA1 and GluA2 subunits as well as NMDA GluN1 and GluN2B subunits were altered in our model. Both GluN1 and GluN2B subunit levels were decreased in SRS animals to 52.8±9.7% (n=6, P<0.05, Fig 5.A) and 68.0±8.5% (n=6, P<0.05, Fig 5.B), respectively, of the total immunoreactivity detected in Sham controls. Likewise, GluA1 and GluA2 levels were similarly decreased in SRSs animals vs Sham controls, showing respectively only 69.8±4.9% (n=6, P<0.05, Fig 5.C) and 77.9±4.5% (n=6, P<0.05, Fig 5.D) of the total Sham immunoreactivity. As a consequence, the GluA1/GluA2 ratio was also smaller (P<0.05, Fig 5.E) in SRS (0.733±0.045, n=6) than in Sham rodents (0.985±0.036, n=6).

**Figure 5.**
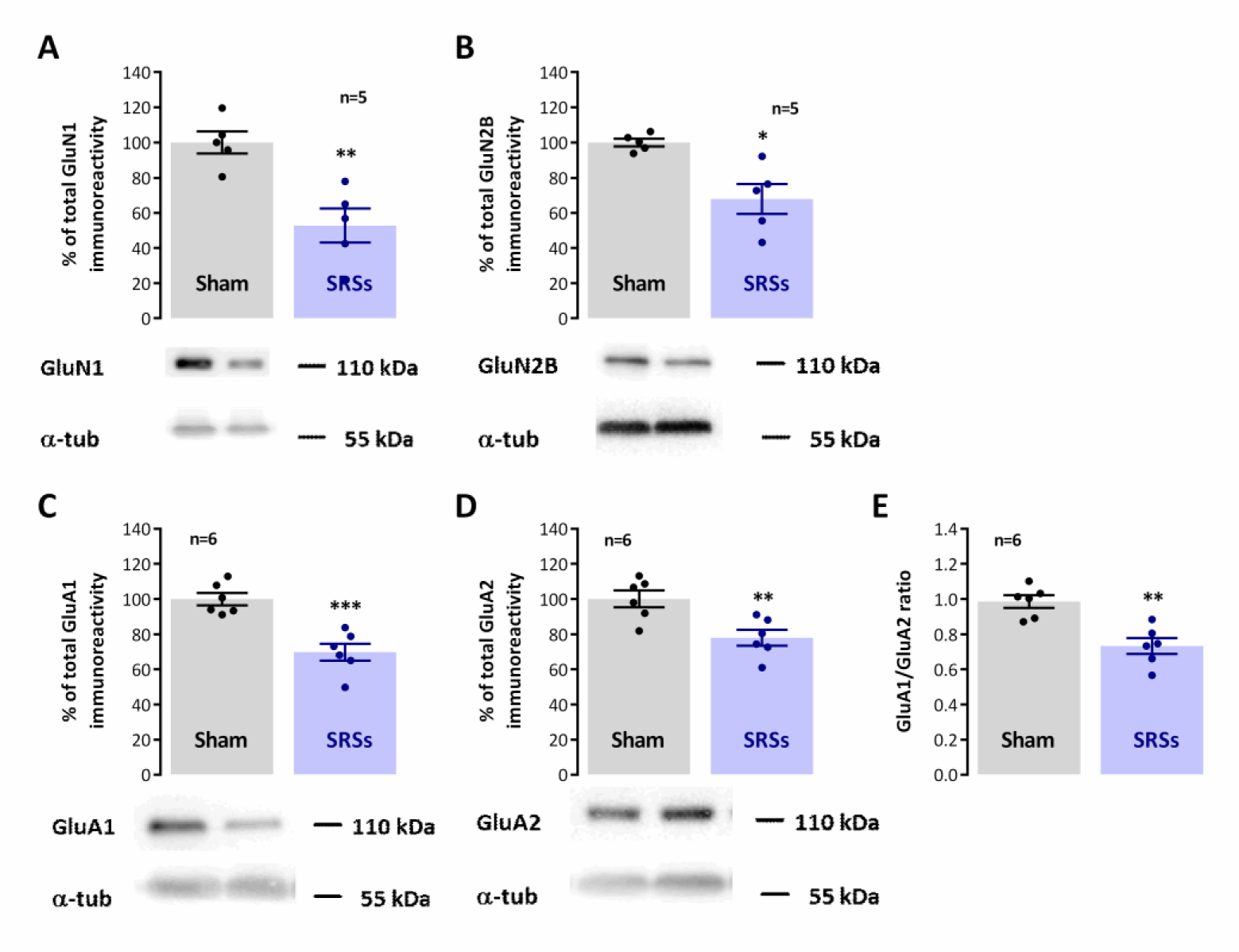
– Hippocampal synaptic AMPA and NMDA receptor subunit composition is altered in the Li^2+^-pilocarpine rat model of epilepsy. Each panel shows at the bottom the western-blot immunodetection of NMDA GluN1 (**A.**) and GluN2B (**B.**) subunits and AMPA GluA1 (**C.**) and GluA2 (**D.**) subunits, obtained in one individual experiment. The GluA1/GluA2 (**E.**) ratio is also depicted Western blot experiments were performed using total hippocampal membranes isolated from six individual animals for both Sham and SRSs. Respective average change NMDA GluN1 (**G.**) and GluN2B (**H.**) together with AMPA GluA1 (**E.**) and GluA2 (**F.**) subunit immunoreactivities are also plotted at the top in each panel. Individual values and the mean ± S.E.M of 5-6 independent experiments are depicted. 100% - averaged target protein immunoreactivity in Sham controls. **\*** P < 0.05 (Student’s t-test) as compared to Sham.

Differences in behavioural parameters, particularly impulsive behaviour, motivation, and depression, in the Li^2+^-pilocarpine model of epilepsy have often been attributed to changes in dopaminergic, noradrenergic, and serotonergic transmission, important modulators of arousal, motivation and attention, all very relevant capacities for both the NOR and MN tasks. As such, we investigated the changes in the levels of enzymes and synaptic transporters associated with the catecholaminergic and serotonergic transmission and correlated them with the levels of constitutive synaptic proteins. In SRSs animals, when compared to Sham controls, we observed a mild decrease in the hippocampal levels of the plasma membrane serotonin (SERT) and dopamine transporters (DAT) to 77.6±6.5% (n=5, P<0.05, Fig 6.A) and 81.9±4.0% (n=6, P<0.05, Fig 6.B) of the observed immunoreactivities in Sham animals, respectively. Conversely, the hippocampal levels of norepinephrine transporters (NET) were increased by 29.6±10.2% (n=5, P<0.05, Fig 6.C) in SRSs vs. Sham rats. The levels of tyrosine hydroxylase, the enzyme catalysing the rate limiting step in the synthesis of catecholamines was also enhanced by 99.8±13.3% (n=6, P<0.05, Fig 5.D) in SRSs vs Sham animals, while the levels of dopamine-β-hydroxylase (DBH), fundamental to the synthesis of catecholamines was increased by 51.9±9.8% (n=4, P<0.05, Fig 5.D).

**Figure 6.**
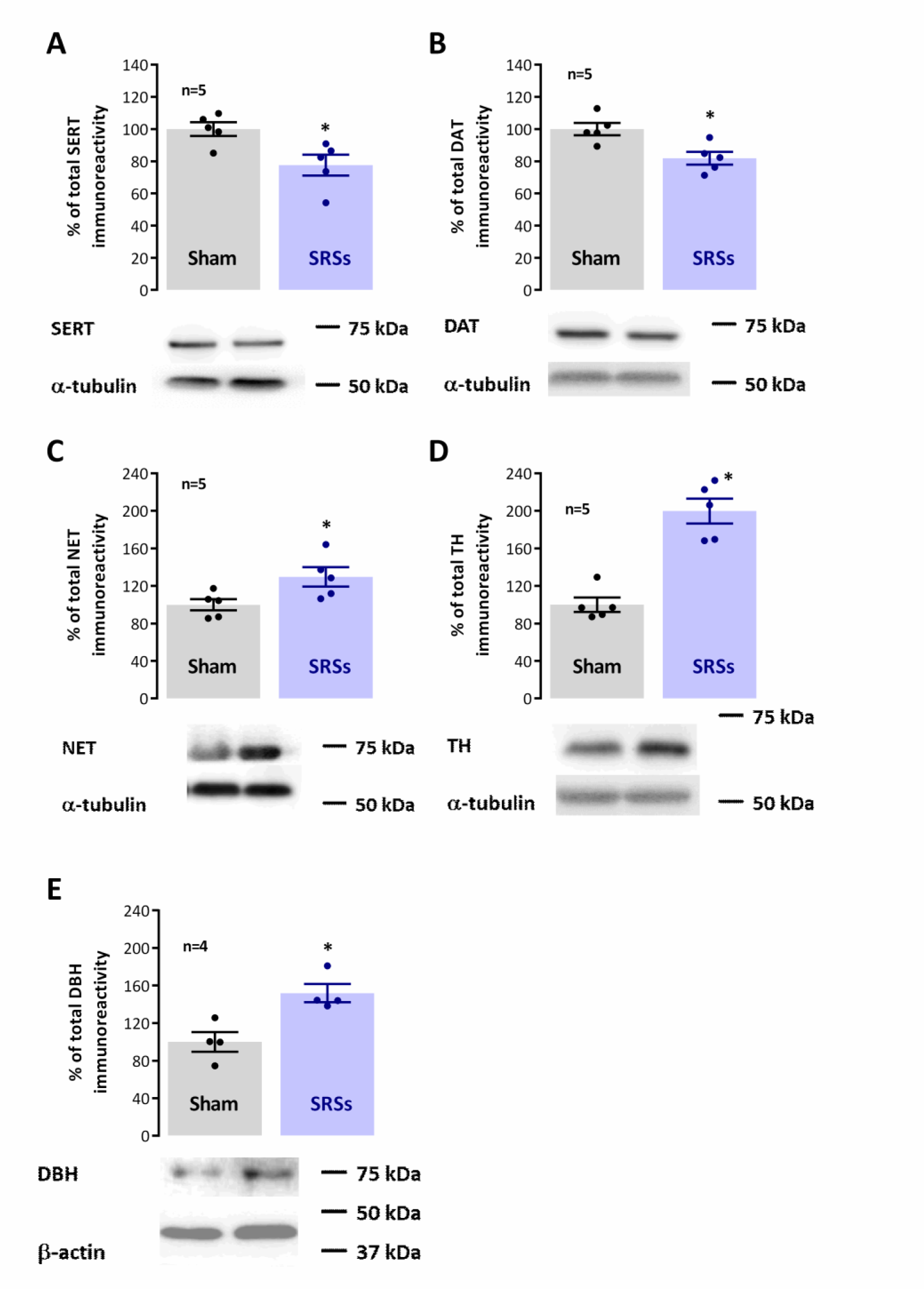
– Hippocampal monoaminergic synaptic content is altered in the Li^2+^-pilocarpine rat model of epilepsy. Each panel shows at the bottom the western-blot immunodetection of SERT (**A.**), DAT (**B.**), NET (**C.**), TH (**D.**) and DBH (**D.**) obtained in one individual experiment. Western blot experiments were performed using total hippocampal membranes isolated from 5-6 individual animals for both Sham and SRSs. Respective average change in SERT (**A.**), DAT (**B.**), NET (**C.**), TH (**D.**) and DBH (**D.**) immunoreactivities are also plotted at the top in each panel. Individual values and the mean ± S.E.M of 4-6 independent experiments are depicted. 100% - averaged target protein immunoreactivity in Sham controls. **\*** P < 0.05 (Student’s t-test) as compared to Sham.

## Discussion

The main findings of the present work are that: 1) exploration of the novel location of known objects in a holeboard is impaired in the Li^2+^-pilocarpine rat model of SRSs; 2) novel object recognition was not significantly altered in SRS animals; 3) the levels of serine and dopamine nerve terminal transporters (SERT and DAT) were mildly decreased in SRS rat hippocampal membranes while 4) the levels of the nerve terminal norepinephrine transporter (NET), of tyrosine hydroxylase (TH) and dopamine-b-hydroxylase (DBH) were enhanced. We also confirmed deficits in spatial learning and alterations in AMPA and NMDA receptor composition and synaptic proteins as found in previous studies in rodent models of epilepsy. Altogether, these observations provide evidence for a disfunction of the novelty processing circuits in SRS animals while characterising monoaminergic transmission disfunction in the Li^2+^-pilocarpine model of SRSs and suggest similar changes this may also occur in human MTLE. As such these could be relevant targets for future pharmacological, behavioural, or possibly combined therapies to mitigate cognitive decline in MTLE.

Cognitive deficits in animal models of epilepsy have extensively been studied, as have the roles of different monoamines in animal models of epilepsy yet previous studies relating cognition and novelty detection impairment to hippocampal monoamines were performed individually, i.e. focusing on one or two neurotransmitters at a time, in distinct animal models of epilepsy, and at different time points following spontaneous recurrent seizures (SRSs) onset, making it difficult to evaluate the relative contribution of each monoamine transmitter to hippocampal-dependent disfunction and its relation to ongoing cognitive disfunction and altered synaptic plasticity. As such, we set out to investigate this in the Li^2+^-pilocarpine model of SRSs.

As mentioned earlier, novelty is an important stimulus in episodic memory formation, and numerous studies have shown that different aspects of novelty have distinct impact on hippocampal dependent learning and synaptic plasticity. The fact that hippocampal long-term potentiation (LTP) or long-term depression (LTD) of synaptic transmission, contribute to encode different aspects of novelty acquisition [11], with LTD being facilitated during the location of new objects or known objects in new locations, and LTP being favoured during exploration of a new environment is by itself proof of a complex interplay of these two forms of synaptic plasticity in hippocampal-dependent cognitive processes. The balance of the two is not only crucial to the formation of a complete spatial map [11,12], but to the consolidation of spatial memory [13,14] and the reversal learning of recently acquired spatial memories [37]. The stability of the memories formed, and of the associated synaptic plasticity phenomena, can be shaped by previously learning experiences (either recent or remote) through metaplasticity [15]. In this respect, novelty, besides being an important trigger for memory acquisition, can also influence ongoing learning and synaptic plasticity events. Spatial novelty enhances retrieval of a previously acquired memory when appearing up to 2h before retrieval, through an NMDA-dependent mechanism [16], while known objects presented in new locations in a familiar environment enhance inhibitory avoidance learning, in a process dependent on hippocampal LTD [13].

New location of known objects in a familiar environment, a behavioural mismatch novelty paradigm, profoundly alters network activity in the CA1 area of the hippocampus in mice [38], and modulates rodent hippocampal synaptic plasticity *in vivo*, through short-term metaplasticity [11]. Mismatch novelty detection, a novelty paradigm remarkably important in memory reformulation and reconsolidation, also triggers the activation of the hippocampal CA1 area in human studies, and is compromised in patients with hippocampal lesions [17,18]. Furthermore, studies from our group showed that repeated exposure to mismatch novelty has a long term metaplastic effect on both LTP and LTD in the hippocampus [19], suggesting that behavioural tasks involving mismatch novelty may be of value in cognitive therapy strategies aiming to mitigate the LTP/LTD imbalance found in aging or diseases like epilepsy or Dowńs syndrome.

In this paper we demonstrate that mismatch novelty detection is specifically compromised in SRS rats as compared to other novelty paradigms like novel object recognition. Altogether, this suggests that recurrent seizures affect more prominently hippocampal neural pathways specifically associated with mismatch novelty detection and processing. Although exploratory responses to open-field exposure were also significantly distinct in SRS and Sham animals, this task, unlike the other two, involves facing an unfamiliar and potentially dangerous environment. As such, amygdala damage also observed in this model [39] may play a role in altered SRS animal performance in this test. Interestingly, performance of SRS rats in the EPM, a test specifically designed to evaluate anxiety traits, revealed that SRS rats are particularly unaware of danger, as previously described [40]. It is believed that altered performance in this test is also related to attention deficits, as the Li^2+^-pilocarpine model of temporal lobe epilepsy has been advanced as a model of attention deficit hyperactivity disorder (ADHD) [36].

Several neurotransmitter systems have been implicated in the hippocampal detection and processing of novelty stimuli. Dopaminergic neurons originating from the VTA and *locus coeruleus* neurons (LC) co-releasing dopamine and norepinephrine innervate the ventral hippocampus, regulate physiological arousal, attention, and motivation and are thought to play an essential role in the efficiency of cognitive function [20,21], playing a crucial role in recognition memory and novelty signalling [23]. Likewise, transmission by *medium raphe* serotonergic fibres and septal cholinergic and GABAergic projections, fundamental for the pacing, engagement and suppression of hippocampal of hippocampal theta rhythm [41–43] and for hippocampal-dependent memory formation [44–46], was shown to be differentially modulated by novelty stimuli [47,48].

Furthermore, altered monoaminergic neurotransmission not only constitutes a risk factor for the development of epilepsy [24] but is either linked to the degeneration or upregulation of ascending projections to the hippocampus and cortex, altered hippocampal monoamine levels or altered distribution and function of monoamine receptors in MTLE patients [25]. Similar observations occurred in experimental models of epilepsy, including (1) selective loss of GABAergic septal projections to the limbic cortex [49], (2) enhancement of cholinergic neurons in the median septum associated with proliferation of hippocampal cholinergic boutons and fibre sprouting [50], (3) loss of limbic-projecting serotonergic neurons from the median raphe (MR) nucleus [26] and decreased serotonin levels [36], and (4) deterioration of dopaminergic projections from the VTA to the *nucleus accumbens* [27] coupled to decreased vesicular monoamine transporter 2 in the temporal cortex and hippocampus [28]. Our observations that in the Li^2+^-pilocarpine model of SRSs hippocampal SERT and DAT levels are decreased, are in agreement with these reports yet partially contradict data showing that SERT immunoreactivity is enhanced in the hippocampus of epileptic patients in risk of SUDEP [51]. Nevertheless, loss of serotonergic and dopaminergic signalling may play a role in the impairment of mismatch novelty processing in SRS rats, as novelty-sensitive dopaminergic neurons in the Human *substantia nigra* have been implicated in declarative memory formation [52]. Furthermore, both 5-HT and dopamine play a role in memory destabilization and reactivation by distinct novelty stimuli, that in turn triggers memory reconsolidation [47,53].

In our work, the levels of TH, DBH and NET were, on the contrary, increased in SRS animals vs. Sham controls. This is overall conflicting with previous knowledge that noradrenaline levels are decreased in the Li^2+^-pilocarpine model, and that this is related to depression-like symptoms [36]. However, these findings are in line with several studies demonstrating that inactivation of NET ameliorates seizures in animal models of epilepsy [54] and with reports that TH and NET are upregulated following seizures in animal models [55,56]. Altogether, this suggests that the ADHD-like phenotype observed in this animal model may be related to an excessive, rather than impaired, noradrenergic signalling. Alternatively, the impairment in serotonergic signalling that is concomitantly observed [36,56] may be the determinant factor in this respect. This question should be further investigated but is currently beyond the scope of this paper.

In this study we also observed a decline in gephyrin levels and PSD-95 levels, specific markers of GABAergic and glutamatergic synapses, that is suggestive of a global decline in both intrahippocampal and external hippocampal-projecting glutamatergic and GABAergic fibres or respective nerve terminals. Since this effect is much more pronounced for gephyrin than for PSD-95 it is to be considered that seizures affect GABAergic transmission more strongly than glutamatergic, at least at this time point after SRS onset. Although this may reflect in part the above-mentioned selective loss of GABAergic septal projections [49] it is also long known that hippocampal GABAergic interneurons involved in disinhibition are particularly affected by seizures and epileptic state [4,57–60], a factor that is determinant in gradually enhanced hippocampal excitability and progressive epileptogenesis. The decline in PSD-95 levels in SRS hippocampal membranes is consistent with previous observations in the kainic acid model of SRSs [61]. This study also describes a concomitant decrease in GluN2B subunits, that is consistent with our current paper, and was expected given the role of PSD-95 in membrane anchoring of NMDA receptor subunits [62,63]. Interestingly, the decline in GluN1 subunits in SRS rats was much stronger, suggesting a major role for these receptors in hippocampal-dependent cognitive decline and epilepsy pathology, and in line with what was observed in previous studies [62,64,65]. Regardless of the multiple controversies generated by studies in different models of epilepsy, at multiple time points following SRS onset and, using multiple approaches to detect levels of synaptic proteins, the role of synaptic reshaping in temporal lobe epilepsy onset and progression is consensual. AMPA and NMDA receptor remodelling plays a crucial role in this process, and altered synaptic plasticity and transmission is a major hallmark of epilepsy animal models but it has been demonstrated also in the human brain [65–67]. Our work confirms in our model that GluA1 and GluA2 levels as well as the GluA1/GluA2 ratio are diminished in SRS animals, as previously described [40], a necessary confirmation given the conflicting results found in the literature.

In conclusion, we investigated the response to mismatch novelty in the Li^2+^-pilocarpine rat model of TLE and its correlation to hippocampal monoaminergic and synaptic markers and other hippocampal dependent learning and memory tasks. We observed an impairment in the exploration of a known environment containing familiar objects presented in a new location in rats showing spontaneous recurrent seizures (SRSs) for at least 4 weeks, suggesting that deficits in mismatch novelty detection indeed contribute to cognitive impairment in MTLE. This was correlated with alterations in the hippocampal monoaminergic system that may contribute to the attention deficit-like profile previously observed in the Li^2+^-pilocarpine model of epilepsy.

## Acknowledgements

The authors thank Institute of Physiology, Faculty of Medicine, University of Lisbon for animal housing and Marta Bento and André Serpa for technical contribution.

---

**Supplementary Figure 1.**
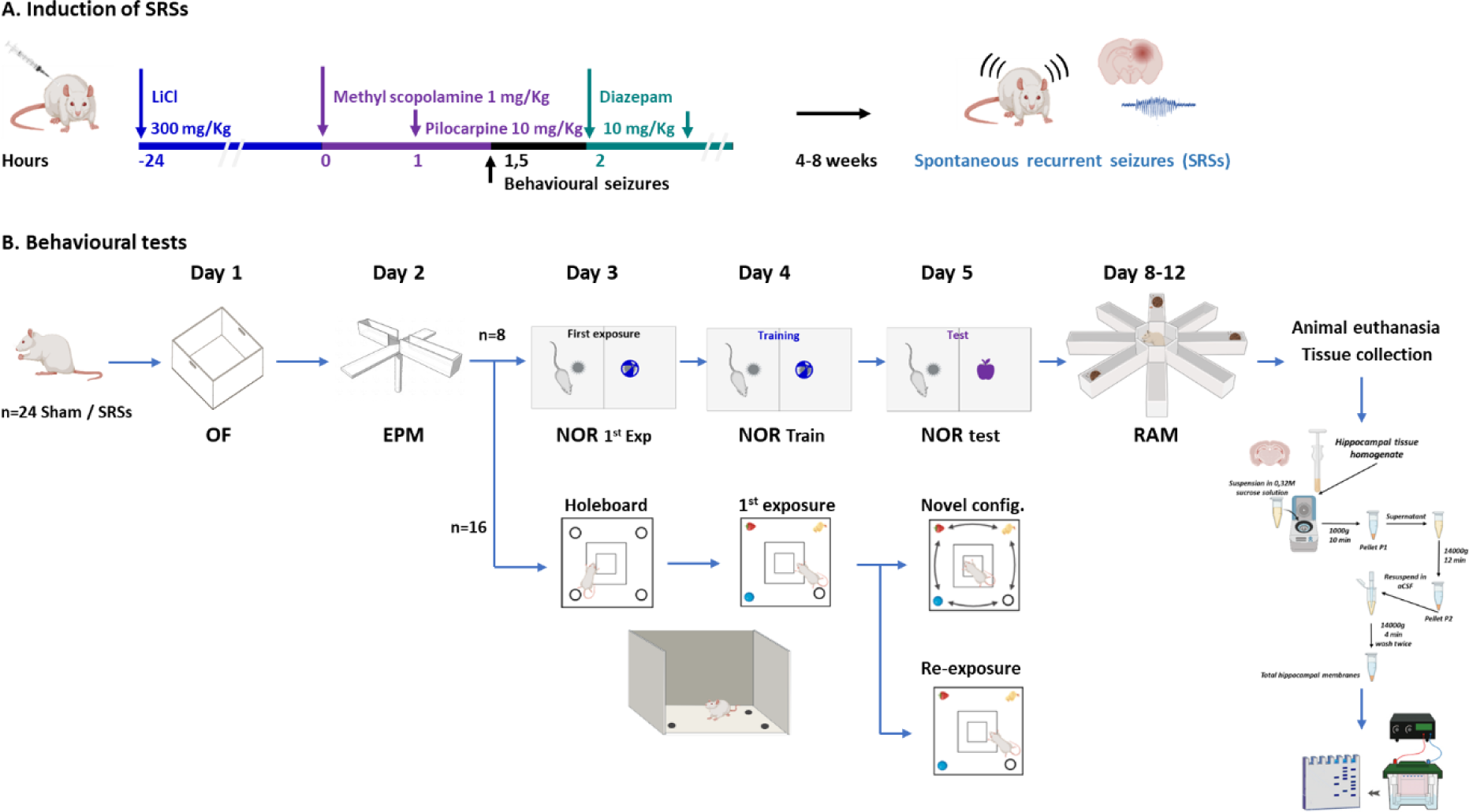
– Experimental workflow for induction of SRSs with Li^2+^-pilocarpine in the rat (A.) and for behavioural evaluation (B.). In A., the sequential drug treatments required to induce and terminate behavioural seizures are shown in the left and the development of spontaneous recurrent seizures (SRSs) within 4-8 weeks are shown on the right. In B., the sequence and division of animals to the different tests is shown sequentially. EPM – elevated plus maze test; NOR – Novel Object recognition test; OF-Open field test; RAM – radial arm maze test for spatial memory.

**Supplementary Figure 2.**
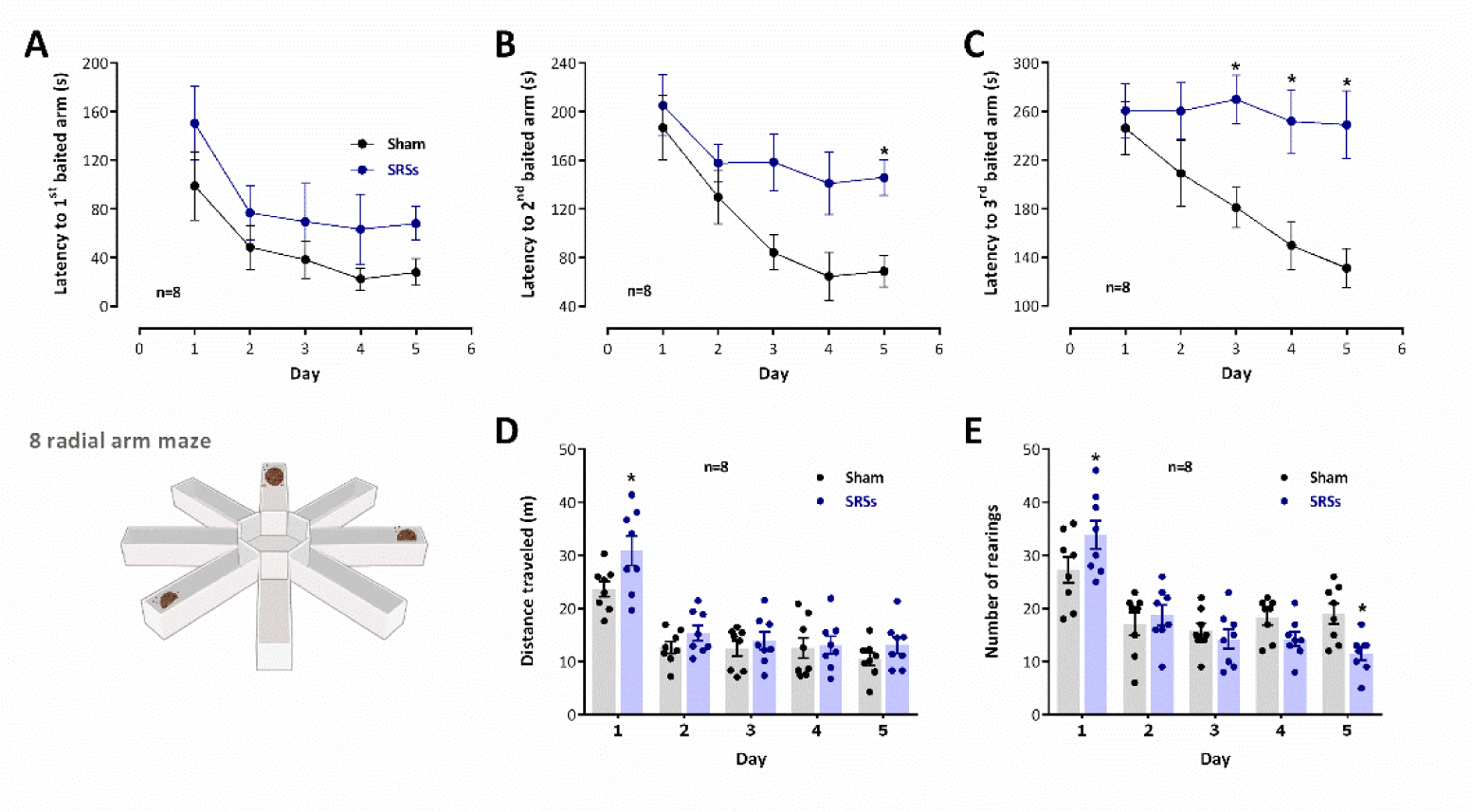
– Impaired learning in the radial arm maze in the Li^2+^-pilocarpine rat model of epilepsy. Learning performance in the RAM was evaluated by the latencies to find the first (**A.**), second (**B.**) and third (**C.**), baited arms. Global exploratory activity in the RAM was evaluated by the total distance travelled (**D.**) and number of rearings (**E.**) during each trial. Total trial duration was of 5 min for each session. A schematic representation of the RAM apparatus is depicted (bottom, left) showing example of one possible configuration for the three baited arms. Values are the mean ± S.E.M. *p < 0.05 vs Sham controls (Two-way ANOVA).

